# Serial amplification of tau filaments using Alzheimer’s brain homogenates and C322A or C322S recombinant tau

**DOI:** 10.1101/2025.03.29.646137

**Authors:** Alessia Santambrogio, Sofia Lövestam, Michael A. Metrick, Thomas S. Lohr, Peifeng Xu, Byron Caughey, Sjors H. W. Scheres, Michele Vendruscolo

**Author notes:** Equal contributions (Alessia Santambrogio, Michael A. Metrick).

## Abstract

The assembly of tau into amyloid filaments is a defining characteristic of Alzheimer’s disease (AD) and other tauopathies. Cryo-electron microscopy (cryo-EM) showed that specific tau folds characterise different diseases, and that *in vitro* models often yield filaments with folds that do not replicate those that form in disease. Here, we investigated the aggregation of full-length recombinant 0N3R tau, using wild-type, or mutations C322A or C322S and a real-time quaking-induced conversion (RT-QuIC) assay with brain homogenate seeding. The assembly of C322A 0N3R tau resulted in filaments with a structure resembling the AD fold in paired helical filaments (PHFs), but with a more open C-shaped core attributed to the C322A mutation. C322S 0N3R tau formed structurally more distinct filaments with respect to PHFs, with an ordered carboxy-terminal region. Both mutant filaments retained the ability to seed a second round of aggregation. Meanwhile, wild-type 0N3R tau exhibited poor reproducibility and formed predominantly unfolded aggregates. Our findings emphasise the need for optimised assembly conditions to obtain disease-relevant filament folds. Refining these methodologies could enhance our understanding of the molecular origins of tauopathies and facilitate the development of targeted therapeutic strategies for these conditions.

## Introduction

The aggregation of microtubule-associated protein tau is a hallmark of Alzheimer’s disease (AD) and a family of related disorders known as tauopathies [1-4]. By the differential splicing of *MAPT* mRNA, six isoforms of tau are commonly found in the adult human brain, ranging from 352 to 441 amino acids in length [2,5,6]. These isoforms are defined by the presence of either zero, one, or two N-terminal inserts (0N, 1N, 2N), which are encoded by exons 2 and 3. Tau contains up to four microtubule-binding repeats (MTBRs) denoted as R1 to R4. The expression of all four MTBRs yields 4-repeat (4R) tau isoforms. Isoforms of tau with only 3 repeats (3R) occur if exon 10, which encodes for the second of the four MTBRs (R2), is spliced out. The tau isoform composition of the filamentous aggregates enables the classification of different tauopathies, which present unique clinical syndromes and affect distinct brain regions [2,3,5,6], into three classes: 3R tauopathies, 4R tauopathies, and mixed 3R/4R tauopathies [2,4,5].

Cryo-electron microscopy (cryo-EM) of tau filaments from multiple tauopathies revealed that specific tau protofilament folds characterise different diseases, providing a structural basis for the classification of tauopathies [4]. Tau filaments extracted from the brains of AD individuals contain two protofilaments, each with eight cross-β strands folded into a C-shape [7]. Two polymorphs, paired helical filaments (PHFs) and straight filaments (SFs), differing in their inter-protofilament packing, are observed in AD [7]. The two protofilaments pack symmetrically in PHFs, whereas the packing in SFs is asymmetrical. The filament core comprises residues 306-378 of tau, thus spanning R3, R4 and ten amino acids of the carboxy-terminal domain. In contrast, the filament cores from 4R tauopathies include residues from R2, making 3R tau monomers incompatible with their elongation [4,8,9]. The filament core from Pick’s disease, which is a 3R tauopathy, also comprises residues in R1, rendering 4R tau incompatible with propagation of the Pick fold [10].

Although distinct tau folds appear to be conserved between patients with the same disease, factors driving tau to adopt one conformer over another are unknown. This is a problem for developing *in vitro* models to study specific tauopathies, as these models should replicate the tau fold observed in diseased human brains. For example, structural analysis by cryo-EM of full-length tau filaments assembled using heparin, RNA, or phosphoserine, showed that the structures of these filaments differ from those of the filaments extracted from human brains [11-13]. Seeded aggregation experiments using brain extracts from AD individuals or corticobasal degeneration (CBD, a 4R tauopathy) in SH-SY5Y cell cultures also yielded filament with structures that differ from the structures of the respective seeds [14].

The limitations of these models are prompting the development of new methods that aim to reproduce disease-specific tau filament structures *in vitro*. Tau filaments identical to AD PHFs were obtained by the *in vitro* assembly of the truncated tau fragment comprising residues 297-391 (tau297-391), in phosphate buffer upon shaking [12]. The addition of sodium chloride led to tau filaments identical to those found from brains of individuals with chronic traumatic encephalopathy (CTE) [12]. However, tau297-391 filaments lacks the characteristic fuzzy coat observed in diseased brains [7], which arises from the residues of full-length tau that do not form part of the ordered core. A further complication in modelling the aggregation process of tau was illustrated by a recent time-resolved cryo-EM study, which revealed that AD and CTE filaments assemble via polymorphic intermediates *in vitro*, highlighting the dynamic nature of tau filament formation [15].

The strategy of recapitulating *in vitro* the assembly of filaments that resemble AD PHFs using brain-derived seeds to induce the aggregation of recombinant monomeric tau [16-20] relies on the observation that the free energy barrier for elongation is lower than that for the spontaneous generation of new seeds [21,22]. This strategy was adopted in the assembly of wild-type recombinant human α-synuclein in the presence of multiple system atrophy (MSA) seeds [23]. However, the resulting structures of the seeded assemblies differ from those of the seeds [23]. Current efforts to achieve the *in vitro* assembly of full-length tau into AD PHFs explored site-specific phosphomimetic mutations, substituting serine or threonine with aspartate or glutamate. Using this approach, full-length tau with twelve phosphomimetic mutations, referred to as PAD12 tau, was reported to yield AD PHF structures through both nucleation-dependent, as well as (at least two rounds of) seeded assembly with AD brain-derived seeds [19]. In another approach, AD-brain-derived tau filaments were used to seed full-length recombinant tau with four glutamate mutations, faithfully reproducing AD PHF structures. However, the resulting recombinant tau filaments lacked the ability to seed recombinant tau monomers in a second-round amplification [17].

In this work, we adopted a particular type of seeded aggregation method, known as real-time quaking-induced conversion (RT-QuIC) assay, which was initially developed to achieve the ultrasensitive detection of prions [24-26], as well as pathological forms of α-synuclein [27-30] and tau [31-33]. The assay leverages the intrinsic self-propagating properties of amyloid filaments in cerebrospinal fluid (CSF),brain extracts, or other biospecimens, offering a framework for reliable molecular diagnosis of associated neurodegenerative diseases [34,35]. Recently, we adapted the RT-QuIC assay to amplify a 3R tau fragment (K12) into amyloid filaments in the presence of trisodium citrate to identify small-molecule binders to tau filaments [36]. However, the structural characteristics of the filaments propagated in that RT-QuIC reaction remained unknown, as filaments recovered from the assay were clumped together and therefore difficult to solve by cryo-EM.

Here, we also use trisodium citrate to develop an RT-QuIC assay using wild-type recombinant 0N3R tau and two variants, carrying a cysteine-to-alanine (C322A) or cysteine-to-serine (C322S) mutation. We then determine the structures of the resulting filaments by cryo-EM, following the seeded-conversion by AD brain homogenates (BH). We also show that the kinetics of the assembly of the two 0N3R tau variants is highly reproducible, and that they yield filaments capable of seeding a second round of amplification. Although the resulting filaments are not identical to AD PHFs, the protofilament interface of the C322A filaments is conserved with the AD PHF fold. In contrast, wild-type 0N3R tau displayed poor kinetic reproducibility and primarily generated unfolded filaments. Overall, by systematically assessing tau filament assembly and amplification kinetics, we provide a mutational landscape study that highlights how specific mutations influence the filament structure and seeding efficiency of tau. Our findings suggest that additional molecular components and/ post-translational modifications are likely necessary to achieve the faithful replication in vitro of disease-associated tau filament structures.

## Results

### In vitro seeded assembly of 0N3R tau variants

Wild-type 0N3R tau and two 0N3R tau variants (C322A and C322S) were expressed in *E. coli* and purified (**Figure S1** and **Methods**). Seeded filament assembly was achieved through rounds of real-time quaking-induced conversion (RT-QuIC) (**Figure 1**). AD brain homogenates were prepared from the frontal cortex using ice-cold phosphate-buffered saline (PBS) and processed with a bead mill. In Round 1, filaments were assembled by introducing a small amount of AD brain homogenate (0.5 ng tissue equivalent) as seeds into a solution of 30 μM (55 μg) C322A 0N3R tau (**Figure 2A**), C322S 0N3R tau (**Figure 2C**), or wild-type 0N3R tau (**Figure 2E**). The reaction was carried out under orbital shaking at 500 rpm (60 s on, 60 s off) in a buffer containing 250 mM sodium citrate and 10 mM HEPES at pH 7.4 (N=7). As a control, cerebrovascular disease (CVD) brain material, which lacks immunohistochemically detectable tau pathology, was used under the same conditions. In these controls, filament formation was not observed within the assay time frame for C322A 0N3R tau, and it was delayed for wild-type and C322S 0N3R tau. The first-round assembly of C322A 0N3R tau showed an initial lag phase of 40 h, followed by a growth phase lasting 60 h, before reaching a plateau phase. C322S 0N3R tau exhibited a faster aggregation profile, with the plateau phase reached after 40 h. The first round of assembly of wild-type 0N3R tau assembly resulted in stochastic aggregation under the tested conditions (**Figure 1E**). Further attempts to assemble wild-type 0N3R tau using a higher seed concentration (**Figure S2A**), or an increased protein concentration (**Figure S2B**), did not yield reproducible thioflavin T (ThT) fluorescence curves. Given the reproducibility of the kinetic profiles obtained in the first round for the C322A and C322S 0N3R variants, we investigated the ability to seed the assembly of second-generation filaments. In this second amplification step, filaments from the first round (2% relative to monomer, 1.1 μg) were used as seeds to assemble either 30 μM (55 μg) C322A 0N3R tau (**Figure 2B**) or C322S 0N3R tau (**Figure 2D**). We observed a reproducible steady increase increase in ThT fluorescence, although at a faster rate compared to the first round of assembly.

**Figure 1.**
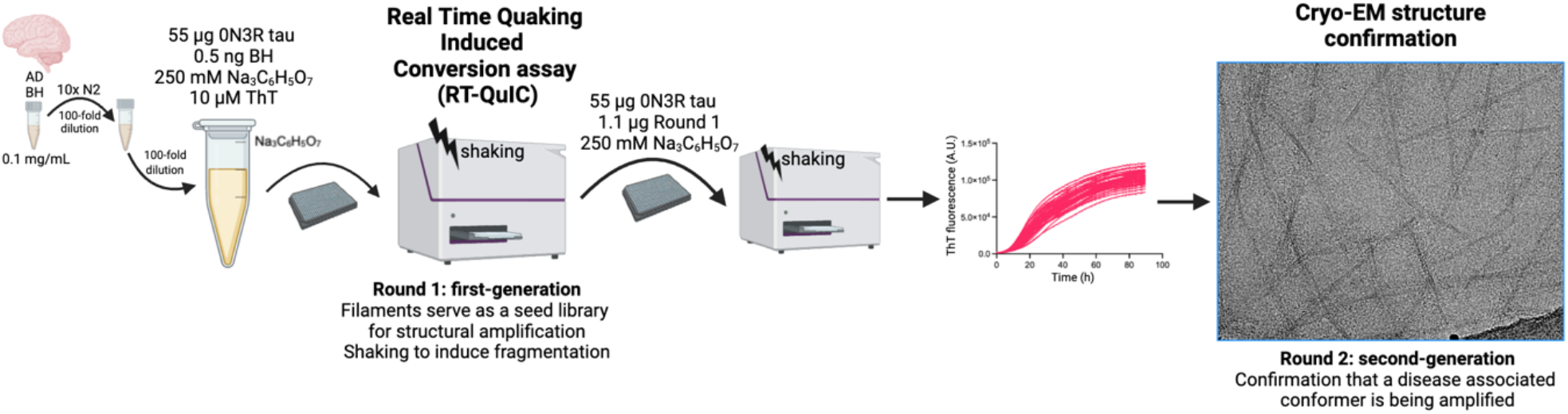
Schematic illustration of the protocol for the two-step seeded assembly of 0N3R tau variants: wild-type, C322A, and C322S. Small amounts of AD brain homogenate (BH) are used for the *in vitro* seeded assembly of recombinant 0N3R C322A/C322S or wild-type tau (first generation, round 1). Amplified filaments are harvested and used as a seed for the second generation of filaments (round 2). AD, Alzheimer’s disease.

**Figure 2.**
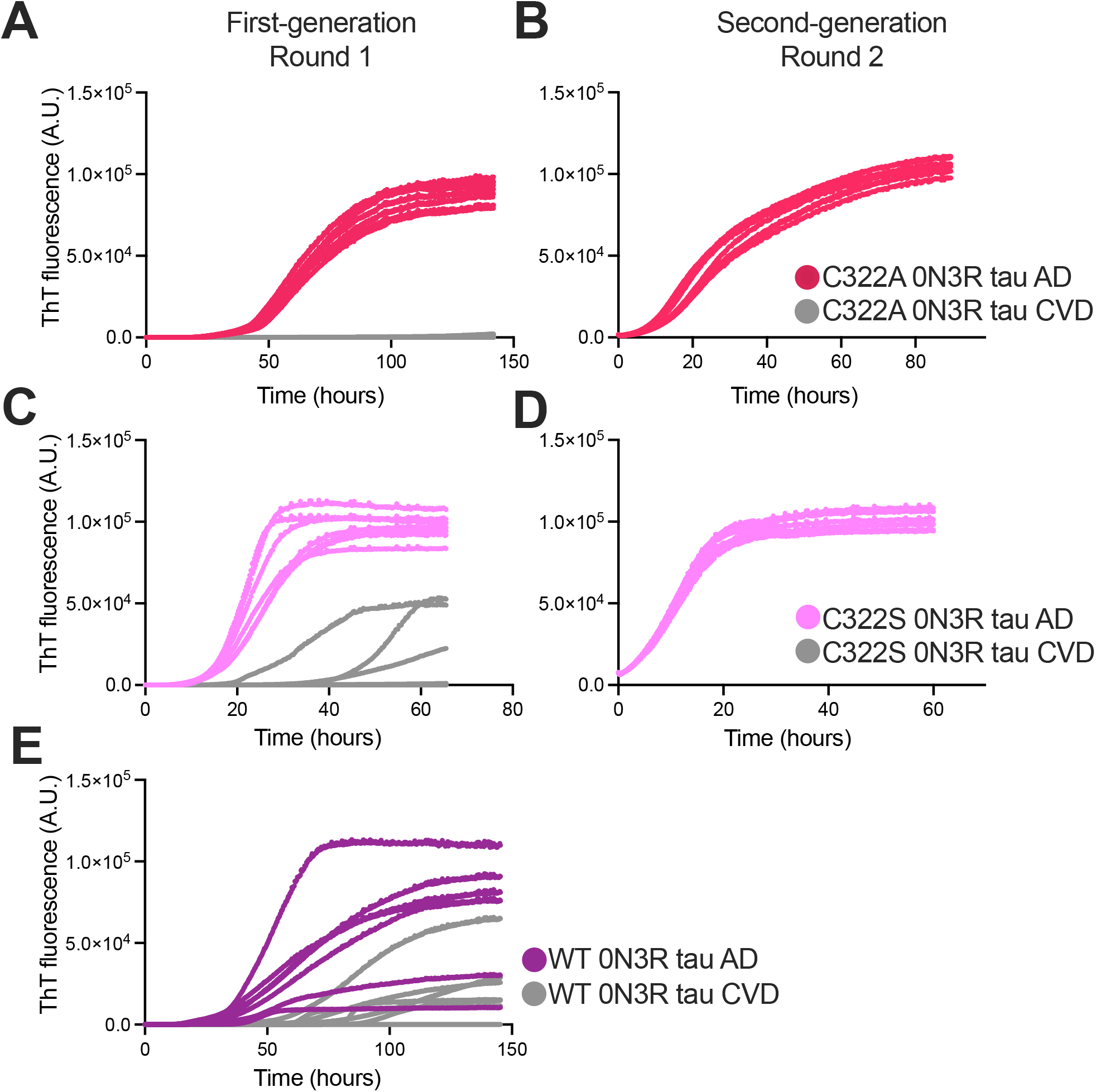
*In vitro* seeded assembly of 0N3R tau. (**A-E**) ThT fluorescence profiles of AD-seeded and CVD-seeded reactions for round 1 (first generation) with C322A 0N3R tau (A), second-round (second generation) seeding (B) with C322A 0N3R tau. Round 1 (C) and round 2 (D) with C322S 0N3R tau, and round 1 (E) with wild-type 0N3R tau N=7. AD, Alzheimer’s disease; CVD, Cerebrovascular disease.

### Cryo-EM structures of 0N3R-C322A and 0N3R-C322S tau filaments after two rounds of seeded assembly

We then determined the cryo-EM structures of the 0N3R-C322A tau filaments after two rounds of seeded assembly. Cryo-EM revealed the presence of multiple filament types, including untwisted filaments, single protofilament filaments, and three types of filaments composed of two protofilaments (**Figure S3**). We solved the structure for one filament type consisting of two protofilaments, as the other filament types were of too low abundance (**Figure S3B**). The structure comprises two C-shaped protofilaments with a fold similar to the AD fold of brain-derived PHFs for residues 323-368 (RMSD 1.9 Å), with the same inter-protofilament packing by residues 331-336 (**Figure 3**). However, at the C322A mutation site the protofilament fold opens up compared to the typical C-shape of the AD fold.

**Figure 3.**
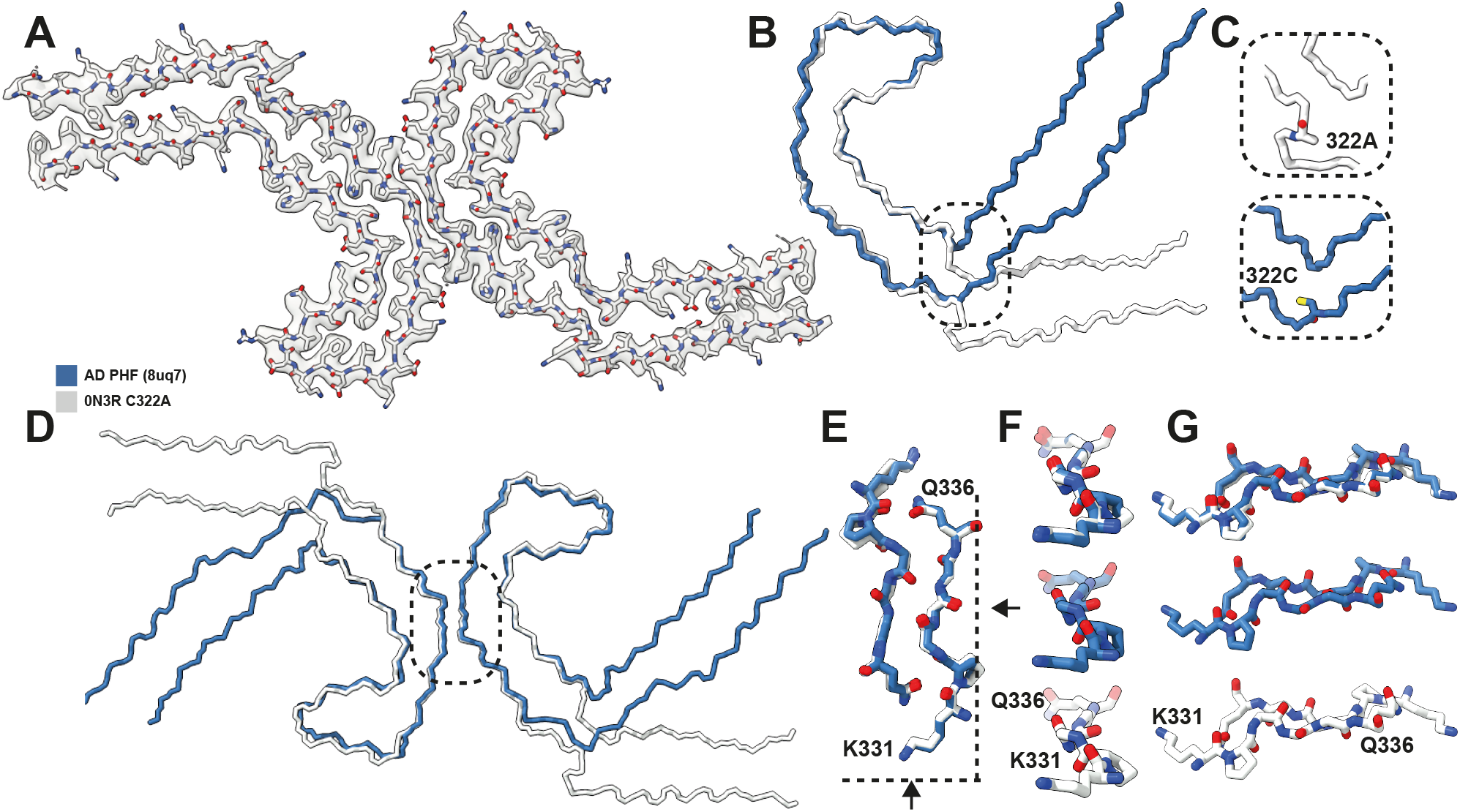
Comparison of the structures of the C322A 0N3R tau filaments and the AD PHF filaments. (**A**) Cryo-EM density map of AD-seeded C322A 0N3R tau filaments (transparent grey) with the all-atom model. (**B**) Comparison of the backbone representation of AD-seeded C322A 0N3R tau filaments (white) and AD PHF filaments (blue) aligned at residues 325-366. (**C**) Close-up at residue 322. **(D)** Comparison of the backbone representations of AD-seeded 0N3R C322A tau filaments (white) overlaid with AD PHF filaments (blue) at residues 332-364. (**E-G**) Close-up of residues 331-336 in different orientations. AD, Alzheimer’s disease.

We also determined the cryo-EM structure of the 0N3R-C322S tau filaments after two rounds of seeded assembly. Cryo-EM revealed the presence of a single type of filament, consisting of one protofilament (**Figure S4**). Although residues 323-378 adopt a conformation that resembles part of the PHF, the C322S mutation results in an outward turn of the amino-terminal part of the ordered core, which only extends to residue S316. Meanwhile, the carboxy-terminal part of the ordered core extends up to residue 434, adopting a three-layer structure, with close contacts between residues 368-380 and 390-411, and residues 415-434 (**Figure 4**).

**Figure 4.**
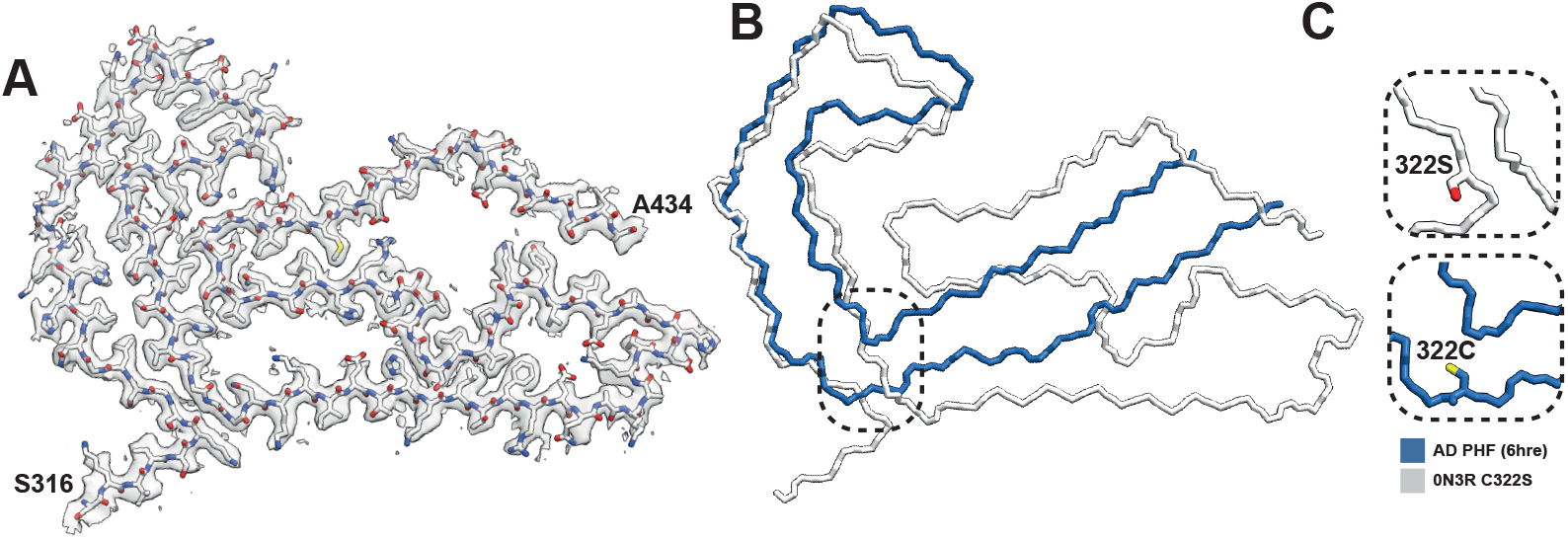
Structure of the C322S 0N3R tau filaments. (**A**) Cryo-EM density map of AD-seeded C322S 0N3R tau filaments (transparent grey) with the atomic model. (**B**) Comparison of the backbone representations of AD-seeded C322S 0N3R tau filaments (white) and AD PHF filaments (blue). (**C**) Close-up of residue 322. AD, Alzheimer’s disease.

## Discussion

The quest for an in vitro assay capable of generating tau filaments with disease-specific folds has been challenging because of filament polymorphism, which implies that assembly of proteins into amyloid filaments may lead to different structures depending on the conditions [3,8,37]. Spontaneous *in vitro* assembly of full-length wild-type recombinant tau into filaments has only been possible with the addition of negatively charged cofactors, such as heparin or RNA [38,39]. However, the resulting structures have been shown to differ from those observed in disease [11,13]. Understanding the factors that influence the formation of distinct amyloid folds is essential for advancing our knowledge of disease mechanisms and ultimately for developing therapeutic interventions.

In this work, to explore the possibility of generating disease-specific tau filaments, we monitored tau aggregation by RT-QuIC in the presence of trisodium citrate [33], focusing on wild-type 0N3R tau and two variants, C322A 0N3R and C322S 0N3R tau. Point mutations that remove cysteine 322 provide an advantage in the purification of 0N3R tau, which can be performed without reducing agents and leads to large amounts of pure protein [33]. The use of tau variants is supported by previous reports that showed that the V337M and R406W familial tau mutation adopt the AD fold, suggesting that certain mutations can be accommodated within the PHF fold [40]. Both the C322A and C322S tau variants exhibit high reproducibility in their aggregation kinetics, as assessed by ThT fluorescence curves, which showed similar half-time and maximum values during the first and second rounds of aggregation (**Figure 2**). Differences in the shapes of the ThT fluorescence curves of the C322S and wild-type reactions, compared to the C322A reaction, may reflect distinct rates of filament fragmentation or elongation.

Two-dimensional class averages of cryo-EM images of the C322A 0N3R filaments after two rounds of assembly showed polymorphism, including single protofilament structures and various types of filaments made of two protofilaments. Our three-dimensional reconstruction of the most abundant polymorph revealed partial templating by AD brain-derived seeds, but the filaments that formed had a more open C-shaped core due to a conformational change at the mutation site. In contrast, the C322S 0N3R variant formed a single filament type comprising one protofilament. Its structure also revealed partial templating by the AD brain-derived seeds, but in this case the ordered core comprised a larger part of the carboxy-terminal terminal domain, extending up to residue 434. This is the third time to our knowledge that it has been reported that a notable portion of the carboxy-terminal becomes ordered in tau filaments, as the assembly of 0N4R tau in the presence of phosphoserine or mouse liver RNA resulted in an ordered core encompassing residues 375–441 and 391–426, respectively [11,12]. Our structures suggest that the residue at position 322 may influence the assembly pathway by modulating protofilament interactions and intermediates, potentially leading to alternative folding trajectories.

We note that it was previously reported that the assembly of full-length wild-type recombinant tau seeded with large amounts of AD brain–derived seeds (10% w/v sarkosyl-insoluble extracts of AD brain) yielded filaments with the AD fold, but the amplified filaments were not able to seed subsequent rounds [17]. Our tau RT-QuIC assay uses considerably lower amounts of AD brain homogenates. We used 0.5 ng of brain tissue per reaction, relative to 55 *μ*g of recombinant tau, representing a more than 10,000-fold diluted seed concentration relative to reactant monomer. However, we observed stochastic aggregation kinetics for the assembly of wild-type 0N3R tau, and the formation of untwisting filaments by cryo-EM. Mutation of cysteine 322 to alanine or serine improved the reproducibility of the aggregation kinetics and led to the formation of filaments that could be used as seeds in a second round of aggregation, but the resulting filaments showed only partial resemblance to AD PHFs. These results suggest that partial templated seeding may occur even with minimal amounts of seeds. Moreover, the ability to propagate full-length tau filaments through multiple subsequent reactions may be useful, as these filaments can serve as a seed library for use in cellular and animal models, or in high-throughput binding assays.

Our findings highlight the importance of aggregation kinetics and structural reproducibility in heparin-free tau RT-QuIC assays, underscoring their potential as a tool for studying templated seeding. The ability to generate seeding-competent filaments of full-length recombinant tau from a small amount of AD brain homogenate suggests that, with further optimisation, RT-QuIC assays could be a useful method for recapitulating disease-relevant structures *in vitro*. For example, the introduction of phosphomimetic mutations in the fuzzy coat, which has been shown to drive the spontaneous assembly of full-length tau into AD PHFs, may also be useful in RT-QuIC assays [19].

Our cryo-EM structures illustrate that even single-point mutations in tau can lead to distinct filament architectures under identical assembly conditions. Thereby, our findings highlight the need for a deeper understanding of the molecular mechanisms that underlie the assembly of amyloid filaments, including the interactions that determine how monomers are recruited into amyloid structures. We anticipate that the identification of the molecular mechanisms that underlie the formation of disease-specific amyloid strains will facilitate their reliable propagation *in vitro*, thus facilitating the development of therapeutic interventions.

## Methods

### Neuropathology and compliance with ethical standards

The procurement and neuropathological analysis of brain samples used in this study were conducted by Bernardino Ghetti at Indiana University School of Medicine. These samples were obtained from deceased, de-identified consenting patients, requiring no further ethical disclosure. In summary, one half of each brain was preserved in formalin, while the other half was frozen. Diagnoses of the fixed tissues were determined using previously established immunohistochemical staining methods [31,41]. Tissue samples were specifically collected from the frontal cortex. Brain homogenates (10% w/v) were prepared from the frontal cortex in ice-cold PBS using 1 mm silica beads (BioSpec, 11079110z) and either a Beadbeater (BioSpec) or a BeadMill 24 (Fischer). The homogenised samples were stored at -80 °C. All sporadic AD (sAD) and cerebrovascular disease (CVD) brain samples listed in Table 1 of Metrick et al. were utilized during the optimisation phase of this study [32]. Following the same sample numbering system, the main figures in this study include data from sAD2 and CVD1.

### Cloning

For the cloning protocol of A233C 0N3R tau and S223 0N3R tau variants, we used the restriction-free (RF) cloning strategy [42]. A233C 0N3R tau and S223 0N3R tau variants were made using restriction-free cloning with wild-type 0N3R tau as the template. Reverse and forward primers were designed to share 14 nucleotides of homologous region and 22 nucleotides for annealing to the template with melting temperatures ranging from 57 °C to 61 °C.

### Protein purification

Wild-type 0N3R tau, C322A 0N3R tau, and C322S 0N3R tau were purified as described previously. Briefly, expression of wild-type 0N3R tau and its variants was carried out in BL21(DE3) *E. coli* as described [22,32]. Cells were grown, and protein expression was induced using an overnight autoinduction method. The cells were harvested by centrifugation (5000 rpm for 25 min at 4 °C) and resuspended in washing buffer (10 mM TRIS, 1 mM EDTA, with or without 10 mM DTT at pH 8.0), supplemented with cOmplete™ EDTA-free protease inhibitor cocktail (10 ml/g of pellet). Cell lysis was performed via sonication (40% amplitude for 5 min, in cycles of 15 s on/45 s off). Lysed cells were slowly poured into washing buffer (2:1, washing buffer: lysed cells) under stirring in a glass beaker, and the temperature was kept above 70 °C. The solution was heated for 10 min and then cooled down using an ice box. Lysates were centrifuged at 14,000× g for 35 min at 4 °C, filtered through 0.45 μm cut-off filters, and loaded onto a carboxymethyl fast flow (CMFF) column. 0N3R tau was eluted from the CMFF resin using a 30-column volume (CV) linear gradient from 0 to 500 mM NaCl. The pooled CMFF eluate was further purified using a sepharose high performance (SPHP) column equilibrated with 10 mM MES, 1 mM EDTA, with or without 10 mM DTT at pH 6.0, and eluted over a 30 CV linear gradient from 0 to 500 mM NaCl. SPHP fractions were pooled, precipitated in acetone, and dissolved in 4 M GuHCl before separation via size-exclusion chromatography (SEC) on a 26 × 600 mm Superdex 75 column equilibrated in 20 mM sodium phosphate, with or without 10 mM DTT at pH 7.4. Purified proteins were lyophilised and stored at -80 °C until further use.

### First-generation seed amplification of 0N3R-C322A tau filaments

AD tau seeds were amplified using 0N3R tau following the method as described previously with several modifications [32]. Sodium fluoride (NaF) was replaced with 250 mM trisodium citrate (Na_3_C_6_H_5_O_7_). 1 μL of each brain homogenate was first diluted to 100-fold with 10x N2 supplement (Gibco). 0.5 μL of diluted brain homogenate solution was again 100-fold diluted to seed 50 μL of first-generation reactions in the presence of 30 μM (55 μg) 0N3R, 10 μM ThT, 250 mM trisodium citrate, 10 mM HEPES, with or without 10 mM DTT at pH 7.4. The final concentration of brain extracts was 0.0001% w/v concentration, equivalent to 0.00001 mg/mL. The total tissue in each 50 μL reaction thus equated to 0.5 ng brain tissue. Reactions were subjected to rounds of 60 s shaking (500 rpm, orbital) and 60 s rest with periodic ThT readings every 15 min at 37 °C in a 384-well Nunc microplate (non-treated polymer base #242764) in a BMG FluoStar lite with aluminium sealing cover to prevent evaporation. Filaments were harvested by scraping and pooling reaction contents once ThT fluorescence reached plateau > 100 h.

### Second-generation 0N3R filament amplification

2% (1.1 μg) of first-generation 0N3R filaments were added to 30 μM (55 μg) monomeric recombinant 0N3R tau. The second-generation reaction buffer contained 250 mM Na_3_C_6_H_5_O_7_ and 10 mM HEPES (pH 7.4), with or without 10 mM DTT. In some reactions, 10 μM ThT was added to enable continuous monitoring of protein assembly, while others, prepared without ThT, were used for cryo-EM analysis. Reactions were again incubated in BMG FluoStar lite at 37 °C with rounds of shaking (500 rpm orbital, 60 s) and rounds of rest (60 s) with periodic ThT reads every 15 min. Filament were recovered for structural analysis by cryo-EM by gently pipetting the reaction solution, avoiding bubbles. Reactions were collected after reaching the ThT fluorescence plateau at > 100 h reaction time.

### Cryo-electron microscopy of 0N3R-C322A tau filaments from RT-QuIC

30 μM 0N3R second generation filaments were recovered from amplification reactions by gentle pipetting. The sample was centrifuged at 2,000 g for 2 min to remover large clumps, diluted by two and 3 μl were applied to glow-discharge discharged R1.2/1.3, 300 mesh carbon Au grids (Quantifoil) that were plunge-frozen in liquid ethane using a Thermo Fisher Scientific Vitrobot Mark IV. Cryo-EM data were acquired on a Titan Krios microscope at the MRC Laboratory of Molecular Biology. All images were recorded at a dose of 40 electrons per Å^2^, using EPU software (Thermo Fisher Scientific). See Tables S1-2 for detailed parameters.

### Helical reconstruction

Movies were gain corrected, aligned and dose weighted using motion correction in RELION [43]. Contrast transfer function (CTF) parameters were estimated using CTFFIND-4.1 [44]. Helical reconstructions were performed in RELION-5.0 [45,46]. Filaments were either picked manually (C322A) or autopicked using Topaz [47,48]. Picked particles were extracted in a box size of 768 pixels and then down-scaled to 128 pixels. Reference-free two-dimensional classification was performed to assess different polymorphs, cross-over distances, and particles for further processing. Initial models were generated de novo from 2D class averages using relion_helix_inimodel2d [49]. Several rounds of 3D auto-refinement was used to optimise the helical twist and rise parameters. Bayesian polishing [43] and CTF refinement [50] were used to further increase the resolution of the final maps. The final maps were sharpened using standard post-processing procedures in RELION. Resolutions were estimated using a threshold of 0.143 in the FSC between two independently refined half-maps [51].

### Model building

Atomic models were built using PDB-ID:6hre as a reference in COOT [52]. Coordinate refinement was performed in ISOLDE [53]. Dihedral angles from the middle rung, which was set as a template, were applied to the rungs above and below to enforce helical symmetry. Separate model refinements were performed on the first half-map using ISOLDE. The resulting model was compared both to the first half-map (FSCwork) and to the second half-map (FSCtest), to confirm the absence of overfitting. Further details of data processing, model refinement and validation are given in Tables S1-2.

### Data availability

There are no restriction on data and materials availability. Cryo-EM maps and atomic models will be deposited in the EMDB and the PDBupon publication.

## Acknowledgements

We acknowledge the expertise of Dr. Heather Greer in the microscopy core of the Department of Chemistry. This work was supported in part by the Division of Intramural Research of the NIIAD, NIH.

