## Supplementary Information for "Serial amplification of tau filaments using Alzheimer’s brain homogenates and C322A or C322S recombinant tau"

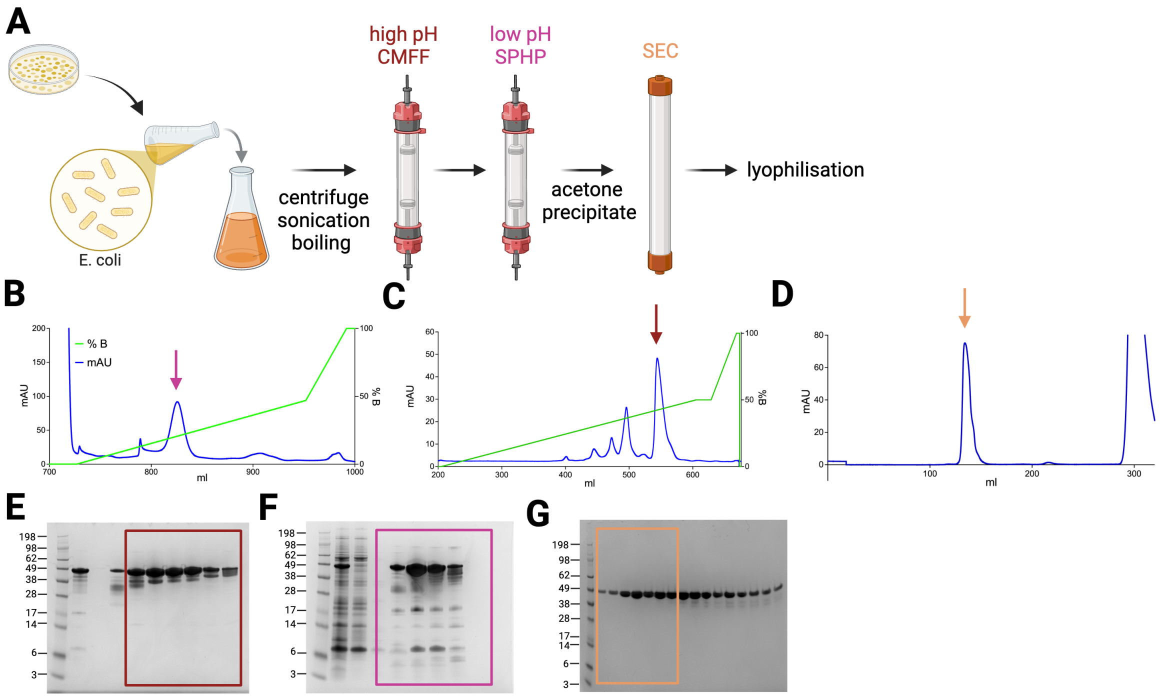

**Supplementary Figure 1. Purification of C322A 0N3R tau**. (**A**) Schematic illustration of the protein purification process, starting from bacterial culture, followed by scale-up, centrifugation, sonication, boiling, CMFF (carboxymethyl fast flow) weak cation exchange chromatography at high pH, SPHP (sulfopropyl high performance) strong cation exchange chromatography at low pH, size exclusion chromatography (SEC), and lyophilization. (**B**) Chromatogram spectra of C322A 0N3R tau via cation exchange chromatography (CMFF) at pH 8 over a linear gradient of 0–500 mM NaCl across 30 column volumes. (**C**) cation exchange chromatography (SPHP) at pH 6 over a linear gradient of 0–500 mM NaCl across 30 column volumes. (**D**) Size exclusion chromatography (SEC). (**E**) Coomassie-stained SDS-PAGE analysis of C322A 0N3R tau after cation exchange chromatography (CMFF) at pH 8. (**F**) SDS-PAGE after cation exchange chromatography (SPHP) at pH 6. (**G**) SDS-PAGE after SEC.

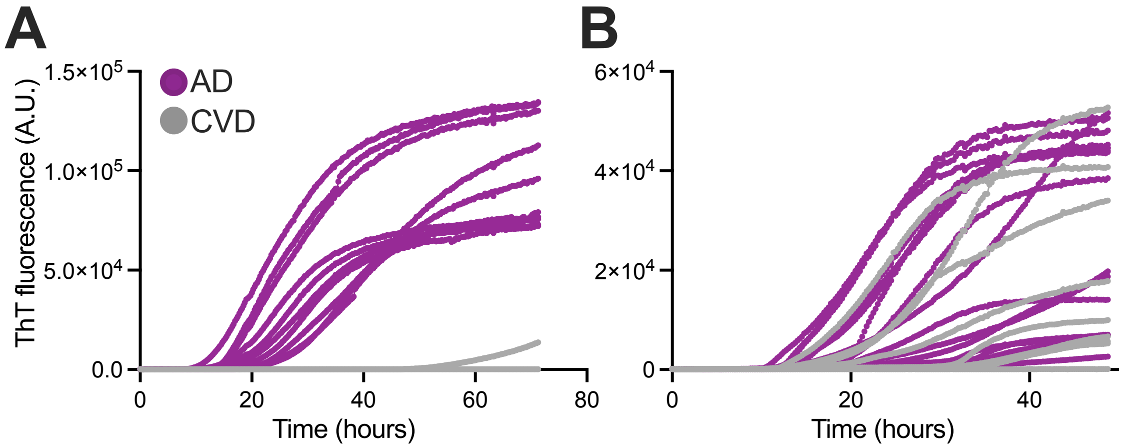

**Supplementary Figure 2. *In vitro* seeded assembly of wild-type 0N3R tau**. (**A,B**) ThT fluorescence profiles of AD-seeded and CVD-seeded reactions for round 1 (first-generation) with C322A 0N3R tau using either a seed concentration of 0.0001% (A) or a higher protein concentration of 50 μM tau (92 μg) (B). N=10; AD, Alzheimer's disease; CVD, cerebrovascular disease.

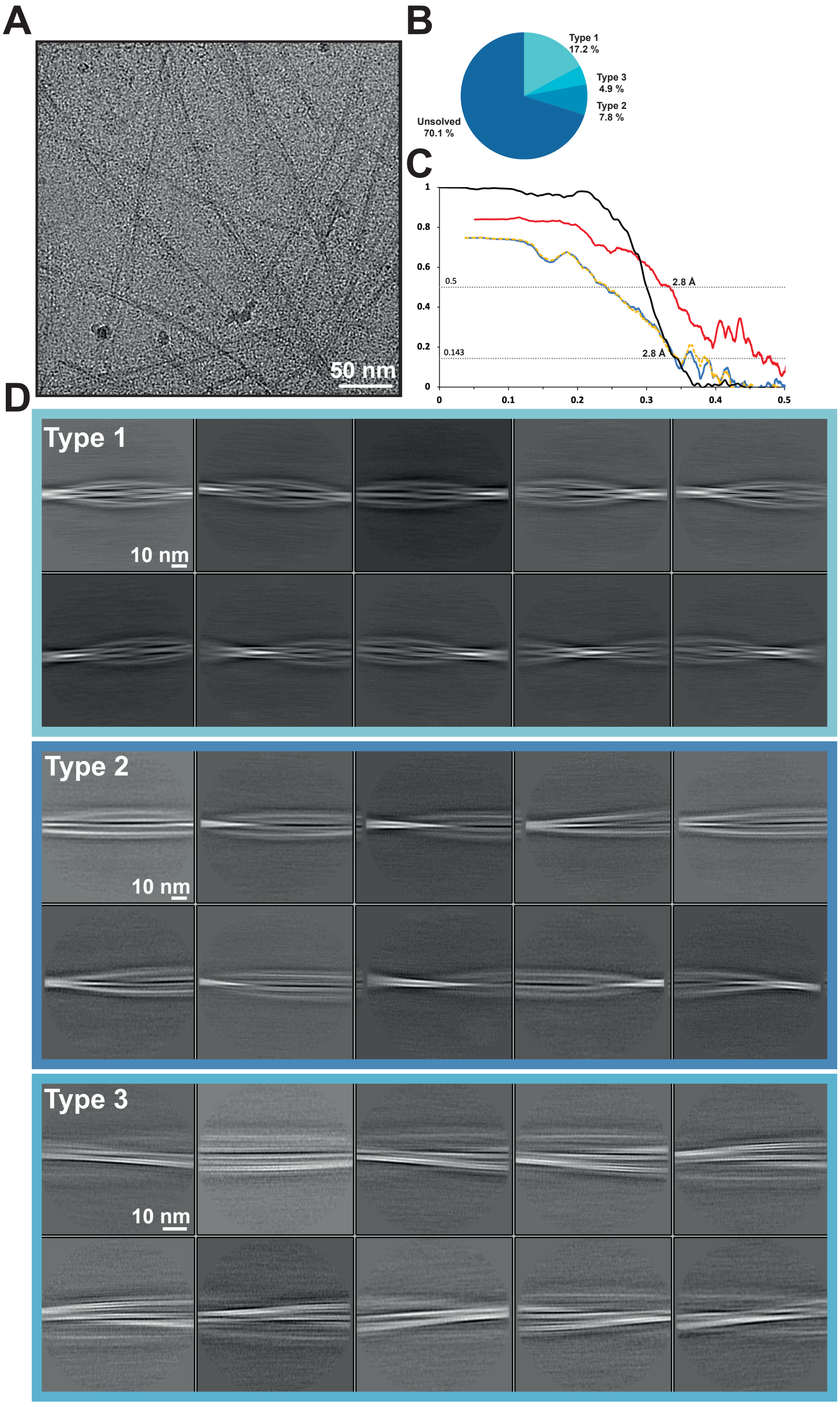

**Supplementary Figure 3. Cryo-EM analysis of AD-seeded C322A 0N3R tau**. (**A**) Cryo-EM micrograph of AD-seeded C322A 0N3R tau. (**B**) Pie chart showing the distribution of filament types. (**C**) Fourier shell correlation (FSC) curves for: two independently refined half maps (black), the final refined atomic model against the cryo-EM map (red), the atomic model refined in the first half-map (blue), and the refined atomic model in the first half-map against the second half-map (yellow). (**D**) Two-dimensional class averages of AD-seeded C322A 0N3R tau filaments.

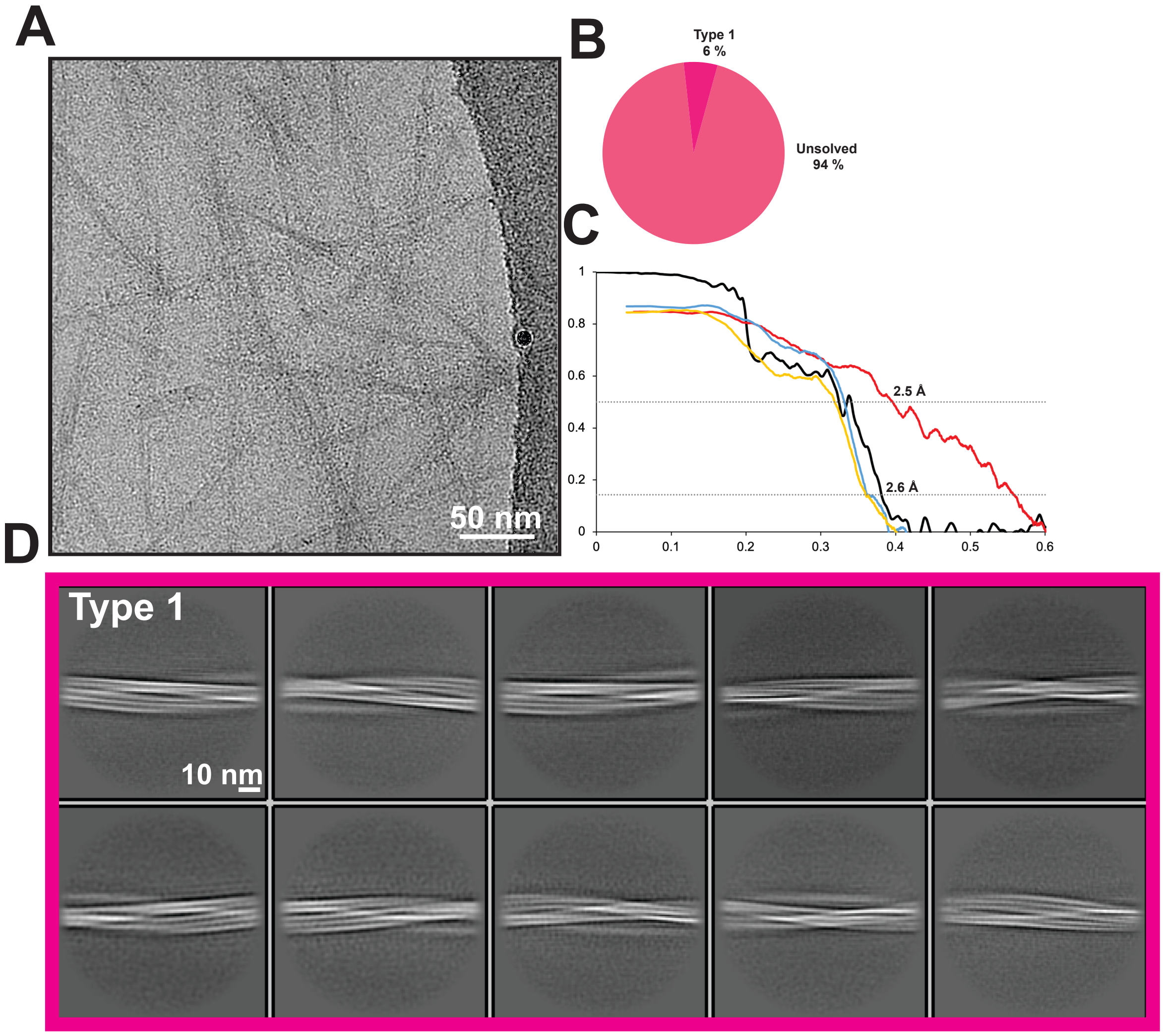

**Supplementary Figure 4**. **Cryo-EM analysis of AD-seeded C322S 0N3R tau**. (**A**) Cryo-EM micrograph of AD-seeded C322S 0N3R tau. (**B**) Pie chart showing the distribution of filament types. (**C**) Fourier shell correlation (FSC) curves for: two independently refined half maps (black), the final refined atomic model against the cryo-EM map (red), the atomic model refined in the first half-map (blue), and the refined atomic model in the first half-map against the second half-map (yellow). (**D**) Two-dimensional class averages of AD-seeded C322S 0N3R tau filaments.

**Table S1: Cryo-EM data statistics and model validation of second-generation 0N3R C322A filaments.**

| **LMB Krios II** | **2nd generation 0N3R C322A** (EMDB xxxx) (PDB xxxx) |
|---|---|
| **Data acquisition** | |
| Electron gun | FEG |
| Detector | Falcon 4i |
| Energy filter slit (eV) | na |
| Magnification | 165,000 |
| Voltage (kV) | 300 |
| Electron dose (e-/Å$^2$) | 40 |
| Defocus range (μM) | 0.5 to 2.5 |
| Pixel size (Å) | 0.824 |
| **Data processing** | |
| Initial particle images (no.) | 516293 (manual) |
| Final particle images (no.) | 61103 |
| Helical twist (°) | 179.599 |
| Helical rise (Å) | 2.417 |
| Symmetry imposed | C1 |
| Map resolution FSC 0.143 (Å) | 2.8 |
| **Refinement** | |
| Initial model used (PDB code) | 6hre |
| Model resolution FSC 0.5 (Å) | 2.8 |
| Map sharpening *B* factor (Å$^2$) | -43.6 |
| Model composition | |
| Non-hydrogen atoms | 3294 |
| Protein residues | 432 |
| Ligands | na |
| *B* factors (Å$^2$) | |
| Protein | 48.4 |
| Ligand | na |
| R.m.s. deviations | |
| Bond lengths (Å) | 0.011 |
| Bond angles (°) | 2.348 |
| Validation | |
| MolProbity score | 1.06 |
| Clashscore | 0 |
| Poor rotamers (%) | 0 |
| Ramachandran plot | |
| Favored (%) | 89.76 |
| Allowed (%) | 10.24 |
| Disallowed (%) | 0 |

**Table S2: Cryo-EM data statistics and model validation of second-generation 0N3R C322S filaments.**

| **LMB Krios III** | **2nd generation 0N3R C322S** (EMDB xxxx) (PDB xxxx) |
|---|---|
| **Data acquisition** | |
| Electron gun | FEG |
| Detector | GATAN K3 |
| Energy filter slit (eV) | 20 |
| Magnification | 165,000 |
| Voltage (kV) | 300 |
| Electron dose (e-/Å$^2$) | 40 |
| Defocus range (µM) | 0.5 to 2.5 |
| Pixel size (Å) | 0.826 |
| **Data processing** | |
| Initial particle images (no.) | 2723519 (autopicked Topaz) |
| Final particle images (no.) | 173842 |
| Helical twist (°) | -0.93 |
| Helical rise (Å) | 4.78 |
| Symmetry imposed | C1 |
| Map resolution FSC 0.143 (Å) | 2.6 |
| **Refinement** | |
| Initial model used (PDB code) | 6hre |
| Model resolution FSC 0.5 (Å) | 2.5 |
| Map sharpening *B* factor (Å$^2$) | -66 |
| Model composition | |
| Non-hydrogen atoms | 2649 |
| Protein residues | 357 |
| Ligands | na |
| *B* factors (Å$^2$) | |
| Protein | 41.4 |
| Ligand | na |
| R.m.s. deviations | |
| Bond lengths (Å) | 0.011 |
| Bond angles (°) | 2.074 |
| Validation | |
| MolProbity score | 0.94 |
| Clashscore | 0 |
| Poor rotamers (%) | 0 |
| Ramachandran plot | |
| Favored (%) | 93.16 |
| Allowed (%) | 6.84 |
| Disallowed (%) | 0 |
